# Bi-modal microwave neuromodulation via thermal and nonthermal mechanisms

**DOI:** 10.1101/2025.03.26.645466

**Authors:** Carolyn Marar, Feiyuan Yu, Guo Chen, Jen-Wei Lin, Chen Yang, Ji-Xin Cheng

## Abstract

Electrical neuromodulation, the current clinical standard, is invasive, expensive, and prone to malfunction. Electromagnetic waves can perform noninvasive neuromodulation, but existing methods are limited by the tradeoff between penetration depth and spatial precision. Microwaves in the 0.9 – 3 GHz range are widely used for telecommunications and can penetrate to the deep brain. Microwaves have been shown to nonthermally modulate neural activity, but the acute bioeffects remain unclear and under-studied. Here, we employ a microwave rod antenna (MRA) to demonstrate bi-modal neuromodulation via thermal and non-thermal mechanisms. The MRA enabled electrophysiological recordings of neurons exposed to microwaves, which elucidated the differential effects of pulsed and continuous microwaves on neurons. These findings build the foundation for developing microwave-based wireless neuromodulation devices for drug-free treatment of seizures and chronic pain.

## Introduction

Nervous system disorders are the leading cause of disability in the world, affecting 3.4 billion people globally1. For conditions involving excessive neural activity, such as epilepsy and chronic pain, medication remains the primary treatment^2,3^. Not only does medication carry the risk of addiction or desensitization, but in many cases, medication is ineffective due to the complex and heterogeneous pathologies of these conditions. Neuromodulation devices present a promising alternative to medication. Neuromodulation is a potentially more robust and precise method for treating neurological disorders by directly altering the firing patterns of target neurons and effecting long-term neuroplastic changes^4^. While neural stimulation has been achieved through various techniques, neural inhibition is less common yet may be more effective at treating conditions involving excessive or synchronized neural firing.

Electromagnetic field and currents have been utilized widely in clinical applications. The current clinical standard for neuromodulation is electrical stimulation, including deep brain stimulation (DBS), vagal nerve stimulation (VNS), and spinal cord stimulation (SCS)^3,5^. Electrical impulses are delivered to target neurons with high spatial precision via an implanted electrode tethered to a subcutaneous stimulator. While electrical stimulation has been used for neural inhibition, its mechanisms remain unclear. Studies suggest electrical inhibition may be achieved via activation of inhibitory networks, depolarization block, or thermal effects^6–8^. While electrical stimulation offers the most precise and reliable neuromodulation, many adverse effects have been reported related to the device and its implantation. Because the stimulator is subcutaneously implanted, any maintenance and repair require surgery and carries the associated risks^9^. Additionally, the complex and tethered design of the device introduces potential for malfunction (cable fracture, lead malfunction, battery failure, etc.). Wireless neuromodulation methods aim to address these issues.

Noninvasive methods, such as transcranial magnetic stimulation (TMS) and transcranial electric stimulation (tES) use external applicators to stimulate deep brain regions. This high penetration depth comes at the expense of spatial precision, increasing the risk of off-target effects^10,11^. Additionally, there is limited evidence for their effectiveness in treating disorders like chronic pain and epilepsy^12,13^. More recently, ultrasound has been explored for neuromodulation^14,15^. Wireless implementation via transcranial focused ultrasound (tFUS) suffers from major interference at the skull, limiting precision and penetration depth.

Microwaves can penetrate over 50 mm into the human brain^24,25^, potentially allowing wireless deep brain neuromodulation. Microwaves, with frequencies of 300 MHz to 300 GHz, have been known to nonthermally inhibit neural activity for decades^16–18^, but the exact mechanisms remain unclear. The cellular mechanisms of microwave neuromodulation are essential to understand due to the ubiquitous use of microwave frequencies, particularly 0.9 – 3 GHz, in telecommunications^19^. Existing studies on the bioeffects of microwaves either focus on high-frequency microwaves (>30 GHz)^18,20^, or on the long-term effects of microwave exposure^19^. There is an alarming lack of studies on the acute cellular effects of 0.9 – 3 GHz microwaves on mammalian neurons. This is largely due to the interference of microwaves with electronics, making it difficult to study fast cellular current dynamics^21^.

Current standards for microwave exposure are primarily based on the known thermal effects, which occur at higher powers^22^. Thus, understanding the nonthermal bioeffects of microwaves and their mechanisms will not only enable more effective use of microwaves for neuromodulation therapy, but also enhance safety standards for all microwave applications. Recently, our team demonstrated wireless, high-precision neural inhibition via a microwave split-ring resonator (SRR)^23^. We further used this device to demonstrate proof-of-concept microwave treatment of epilepsy in an in vivo mouse model. Our data suggested that the neural inhibition is achieved via a non-thermal mechanism. However, the detailed biophysical mechanisms remain to be studied.

In this study, we present the development and application of a miniaturized microwave rod antenna (MRA) to achieve bimodal (non-thermal and thermal) neuromodulation. The MRA can perform neural inhibition as well as induce depolarization via programmed microwaves (pulsed vs. 1 second continuous). Because the MRA concentrates the microwave field similarly to the SRR, we are able to use much lower powers, effectively mitigating electrical artifacts in electrophysiological recordings. This allows us to measure acute changes in neuronal membrane properties. Using a combination of calcium fluorescence imaging and patch clamp recording, we present evidence that neural inhibition occurs through nonthermal perturbation of membrane properties by millisecond pulsed microwaves, while depolarization occurs via induction of thermal transients via 1-second continuous microwaves. In contrast to other neuromodulation techniques which require higher powers to achieve inhibition^6,26^, microwave inhibition occurs at lower powers than depolarization. This study has applications in wireless bimodal neuromodulation to treat disorders like chronic pain and epilepsy. Additionally, this work helps to elucidate the cellular mechanisms of microwave neuromodulation toward the goal of improving safety standards.

## Results

### The microwave rod antenna enables microwave neuromodulation at low dosages

Mechanism studies of microwave neuromodulation are difficult to conduct due to the interference of 0.9-3 GHz microwaves with electronic detectors including, cameras and electrophysiology recorders. To minimize such electrical interference, very low microwave powers must be used, but the cellular effects in this range may not be detectable. We previously demonstrated that 0.66 W/cm^2^ microwave at 2.05 GHz pulsed at 10 Hz for 10 s can inhibit neurons when used with a split-ring resonator (SRR)^23^. However, when the same microwave was applied without the SRR, no significant effect was observed (**Figure 1A, Fig. S1**). Therefore, in order to perform electrophysiology under microwave treatment, an antenna must be used to locally amplify the field to see neuromodulation effects at low powers. To this end, we designed a microwave rod antenna (MRA) with a geometry that is more compatible with recording setups than the SRR. The MRA acts as a dipole antenna, wirelessly coupling the microwave and producing concentrated electric fields at either end (**Figure 1B**). In simulations, a titanium MRA with length 7 mm and diameter 0.5 mm amplified the electric field over 36 times (**Figure 1C**). The length of the MRA is half the wavelength of the microwave in water, meaning its resonant frequency can be tuned by changing its length (**Figure 1D**). The strength of the electric field can also be enhanced by tapering the MRA tip. In simulations, a 7 mm MRA with a 0.5 mm long tapered tip amplified the electric field over 65 times (**Figure 1E-F**). This enhancement, however, comes with a tradeoff between field strength and spatial precision (**Fig. S2**). For this study, we chose a titanium MRA with a 0.5 mm diameter blunt tip so that the field could reach cells within ~1 mm from the tip.

**Figure 1:**
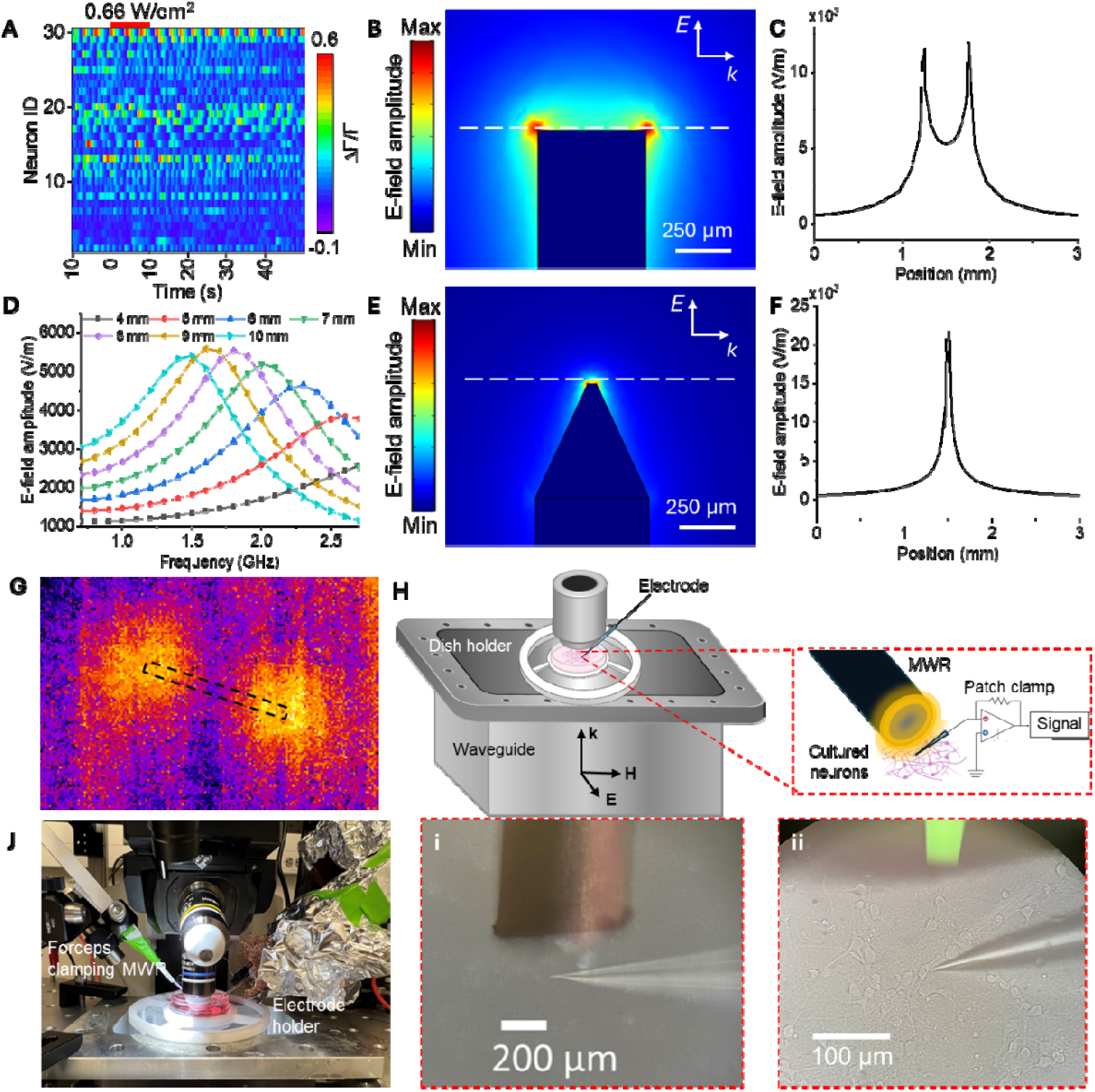
The MRA wirelessly couples and concentrates the microwave field. **(A)** Heat map of calcium fluorescence traces for neurons under 0.66 W/cm^2^ pulsed microwave; **(B)** Simulation of a titanium rod antenn with blunt ends; **(C)** Profile of electric field amplitude at the cross section in B; **(D)** Frequency sweep of electric field amplitude for rod of varying length; **(E)** Simulation of a titanium rod antenna with tapered ends; **(F)** Profile of electric field amplitude at cross section in E; **(G)** Thermal image of blunt rod under a microwave field; outline indicates position of rod; **(H)** Schematic of patch clamp experiments using the microwave rod; **J)** Pictures of the patch clamp setup: i) closeup of rod tip under microscope, ii) closeup of patched neuron next to rod tip.

To validate electric field concentration, the MRA was exposed to continuous microwave and imaged with a thermal camera (**Figure 1G**). Hotspots formed at each end of the MRA with a peak at 2.1 GHz, making this the resonant frequency. Because the RMA amplifies the microwave field at its tips, we were able to perform experiments at power densities as low as 0.066 W/cm^2^. At these power densities, the MRA can be used to perform electrophysiology with minimal electrical interference. For in vitro experiments, the rod was placed over neurons and nearby cells were imaged and/or patched with a glass micropipette (**Figure 1 H-J**).

### The MRA under 10 s microwave pulse train inhibits neurons via a nonthermal mechanism

Previously, our team demonstrated that the microwave split-ring resonator (SRR) inhibits neural activity when the microwave was pulsed at 10 Hz with 10% duty cycle (**Figure 2A**). In the current work, the microwave is pulsed in this manner to minimize the rate of heating and therefore thermal stimulation effects. A 7.5 mm MRA with a 0.5 mm diameter blunt tip was fabricated from pure titanium (**Figure 2B**) and placed over primary rat cortical neurons in vitro. Neurons were dyed with Oregon green to visualize calcium activity. The neurons exhibited spontaneous baseline activity. When the 10 Hz pulsed wave (PW) microwave was applied, the neural activity was strongly inhibited before returning to baseline activity levels (**Figure 1C**). To determine the lower limit of the inhibition effect, microwave power density was decreased from 0.66 W/cm^2^ to 0.066 W/cm^2^ (**Figure 2D-G**). Calcium traces were normalized and baseline-corrected (**Fig. S3**) and area under the curve (AUC) was used to quantify neural activity before, during, and after microwave treatment (**Figure 2H-K**). Activity was decreased by 26.6%, 26.9%, 19.5%, and 3.6% during 0.66 W/cm^2^, 0.41 W/cm^2^, 0.25 W/cm^2^, and 0.066 W/cm^2^ microwave treatment, respectively (N = 3 dishes). 0.066 W/cm^2^ was the lowest power density at which significant inhibition was observed, making it more effective than the SRR at lower power densities (**Fig. S4**). Extended calcium fluorescence recordings confirmed the neurons were not damaged after inhibition (**Fig. S5**).

**Figure 2:**
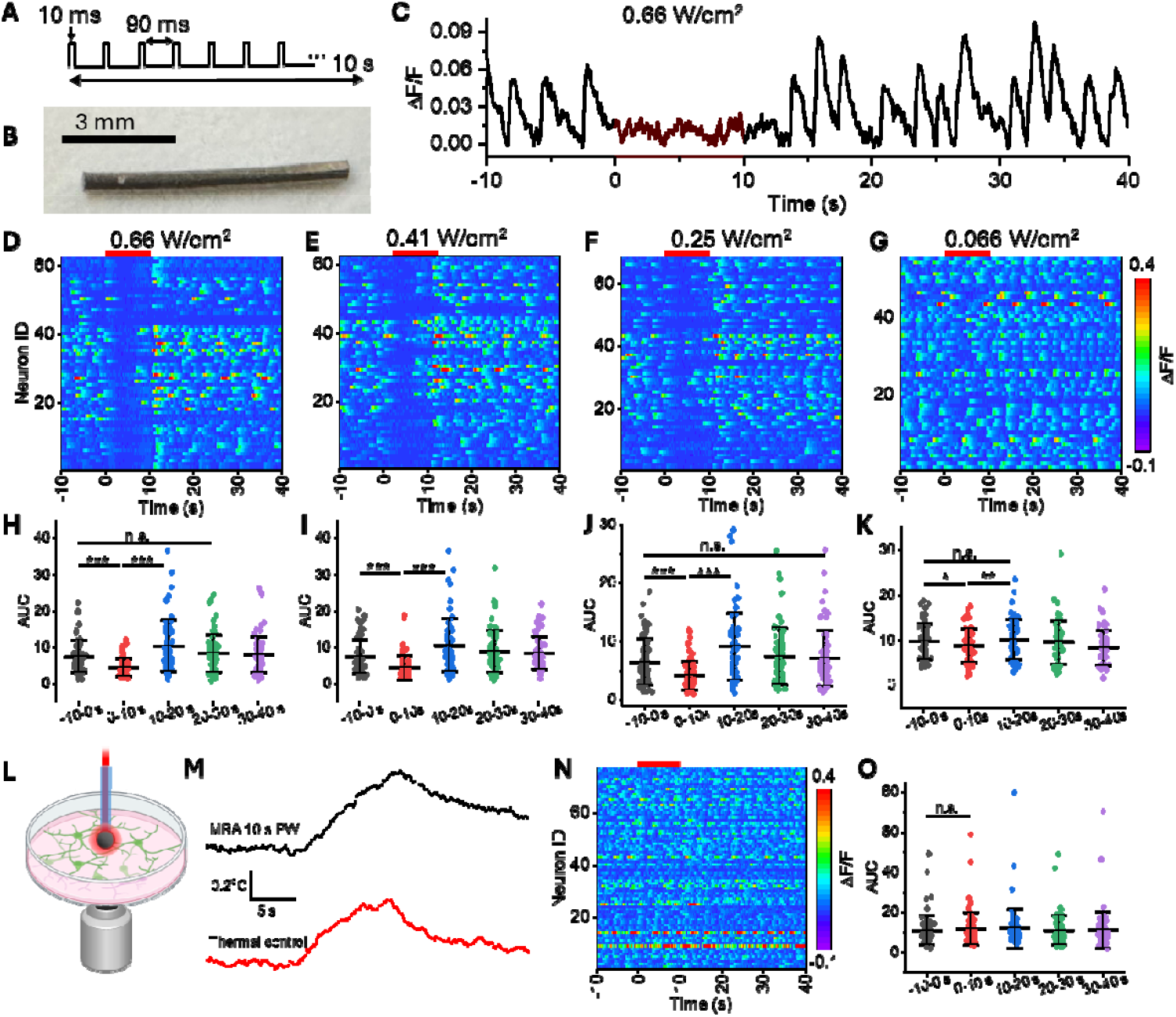
Pulsed microwave inhibits neurons via a nonthermal mechanism. **(A)** Schematic of microwave pulse modulation; **(B)** 7.5 mm blunt tip MRA used in experiments; **(C)** Example trace of calcium fluorescence under 0.66 W/cm^2^ pulsed microwave; **(D-G)** Heat maps of calcium activity in neurons under pulsed microwave at 0.66 – 0.066 W/cm^2^; **(H-K)** Area under the curve of calcium traces for neurons in D-G, respectively; **(L)** Schematic of thermal control experiments; **(M)** Change in temperature near the MRA tip (top) and thermal control trace (bottom); **(N)** Heat maps of calcium activity in neurons under thermal control heating; **(O)** AUC of calcium traces for neurons in N; Plots show mean and standard deviation; Statistical significance calculated using pair sample t test where ***p<0.05, **p<0.01, **p<0.001, n.s. not significant; Red bars indicate treatment periods.

To determine whether the neural inhibition effect of the microwave was thermal, a negative thermal control was performed. The tip of an optical fiber was coated with carbon and heated with a 1064 nm laser to produce heat without microwaves (**Figure 2L**). The laser was pulsed at 10 Hz with 10% duty cycle (same as the PW microwave) to reproduce the temperature profile as closely as possible (**Figure 2M**). Under 0.66 W/cm^2^ PW, the MRA produced a 0.6°C change. For the negative thermal control, the temperature change was 0.53°C with a similar temperature profile. When heated in the absence of microwave, no significant change in neural activity was observed (N = 3 dishes) (**Figure 2N-O, Fig. S6**).

### The MRA under 1 s continuous microwave depolarizes neurons via a thermal mechanism

To determine how pulse modulation and thermal profile impact microwave neuromodulation, the MRA was placed over neurons in vitro (**Figure 3A**) and continuous (CW) microwave was delivered for 1 s. Techniques like infrared neural stimulation induce temporal temperature gradients (dT/dt) to depolarize neurons^27^. Therefore, 1 s CW was used because it is the same total energy dosage as the 10 s PW condition but generates a much higher dT/dt. When 1 s CW 0.66 W/cm^2^ microwave was delivered, neurons exhibited an increase in calcium fluorescence (**Figure 3B**). The microwave-stimulated peak, however, appeared to be a subthreshold depolarization. In contrast to the rapid rise time of spontaneous calcium peaks (blue arrow), the stimulated peak increased slowly until the microwave turned off, after which it decreased. This depolarization is likely due to an influx of Ca^2+^ current since an action potential was not induced. This depolarization was repeatable and non-toxic to the neurons (**Fig. S7**).

**Figure 3:**
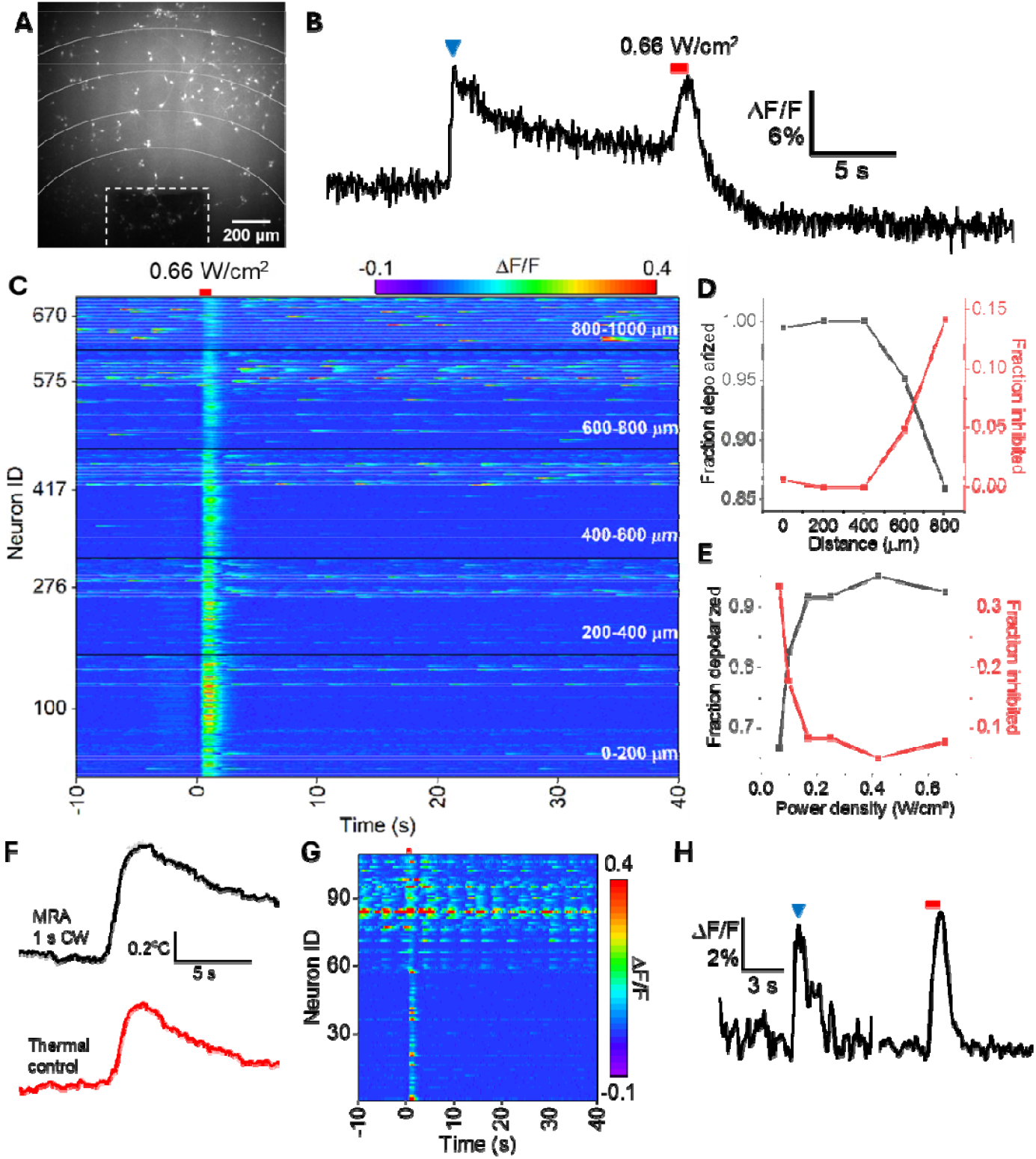
Continuous microwave depolarizes neurons via a thermal mechanism. **(A)** Oregon Green fluorescence image of neurons near MRA tip (dashed outline); **(B)** Example trace of a neuron with spontaneous (blue arrow) and microwave-stimulated (red bar) peaks; **(C)** Heat map of fluorescence traces for neurons at varying distance from the MRA tip under 1 s continuous microwave treatment; **(D-E)** Fraction of cells depolarized and inhibited as function of distance from the MRA tip (D) and microwave power density (E); **(F)** Temperature change traces for the RMA under 0.66 W/cm^2^ 1 s CW microwave (top) and recreated trace using laser heating; **(G)** Heat map of fluorescence traces for neurons at under laser heating; **(H)** Example trace of spontaneous (blue arrow) and heat stimulated (red bar) calcium peaks.

To investigate the distance-dependence of the MRA-induced depolarization, cells were grouped by distance from the MRA tip (200 μm intervals) (**Figure 3C**). As distance increased, the amplitude of the depolarization appeared to decrease. More interestingly, some cells seemed to be inhibited at further distances from the MRA. Cells were classified into two groups based on whether the AUC increased (depolarized) or decreased (inhibited) during microwave treatment (**Figure 3D**). This analysis revealed that the fraction of cells depolarized decreased with distance, while the fraction of cells inhibited increased. This observation is consistent with a nonthermal mechanism for inhibition, as cells further from the MRA experience less heating, but may still be inhibited by a distinct mechanism. As further evidence, depolarization and inhibition fractions were measured at power densities ranging from 0.066 – 0.66 W/cm^2^ (**Figure 3E**). As expected, as power density decreased, depolarization decreased while inhibition increased.

To confirm that microwave-induced depolarization is thermal, a positive thermal control was performed. As previously described, a carbon-coated optical fiber was used to reproduce the thermal profile of the 1 s CW microwave on the MRA (**Figure 3F**). The MRA produced a temperature increase of 0.78°C and the reproduced thermal profile had an increase of 0.60°C. When the cells were heated by the carbon-coated fiber for 1 s, the neurons were depolarized similarly to the 1 s CW microwave condition (**Figure 3G**). Just as in the microwave case, the stimulated peak had a slower rise time than the spontaneous calcium peaks (**Figure 3H**, blue arrow). As expected, microwave depolarization seems to be dependent on dT/dt. Although 0.66 W/cm^**2**^ 10 s PW has a higher absolute temperature increase than 0.41 W/cm 1 s CW^2^ (**Fig. S8**), the former inhibits neurons while the latter depolarizes neurons. This is likely because dT/dt for the CW case is higher than the PW case. These results suggest that microwave depolarization occurs through thermal mechanisms, consistent with the photothermal stimulation techniques.

### Real-time patch clamp recording reveals differential effects of continuous and pulsed microwave on neuronal membranes

To elucidate the distinct thermal and nonthermal effects of microwave neuromodulation, the MRA was integrated into a patch clamp recording system as shown in **Figure 1J**. With the MRA, microwave effects were achieved at lower operational powers, reducing electrical artifacts (**Fig. S9A**). The MRA was held over the neurons and whole cell patch clamp recordings were performed on cells within 1 mm of the tip to measure changes in passive membrane properties and action potential (AP) characteristics. Positive current was injected into the cell every 1 s to induce an AP.

When neurons were exposed to 0.25-0.66 W/cm^2^ 1 s CW microwave, a subthreshold membrane depolarization was observed (**Figure 4A**). The depolarization gradually increased throughout the duration of the microwave, similarly to the calcium imaging data, further supporting a thermally induced mechanism. In some cases, AP amplitude decreased slightly during microwave (**Figure 4Bi**), but there were no other notable changes to AP characteristics (**Figure 4C, Fig. S9B, Fig. S10**). The decrease in AP amplitude can be explained by the microwave-induced depolarization, as there was a correlation between A/A_0_ and ΔV_rest_ (**Fig. S11A**). This indicates that the change in AP amplitude is likely due to inactivation of voltage-gated sodium channels in the cell. Negative current injection revealed a slight decrease in membrane resistance (R_m_) and a significant decrease in membrane capacitance (C_m_) (**Figure 4Bi-C**). Both parameters recovered after microwave exposure. In combination with the calcium influx observed during calcium imaging, this data suggests that the depolarization may be induced by nonspecific cationic current. Since it is thermally induced, the current likely comes from thermosensitive transient receptor potential (TRP) channels, which are nonspecific.

**Figure 4:**
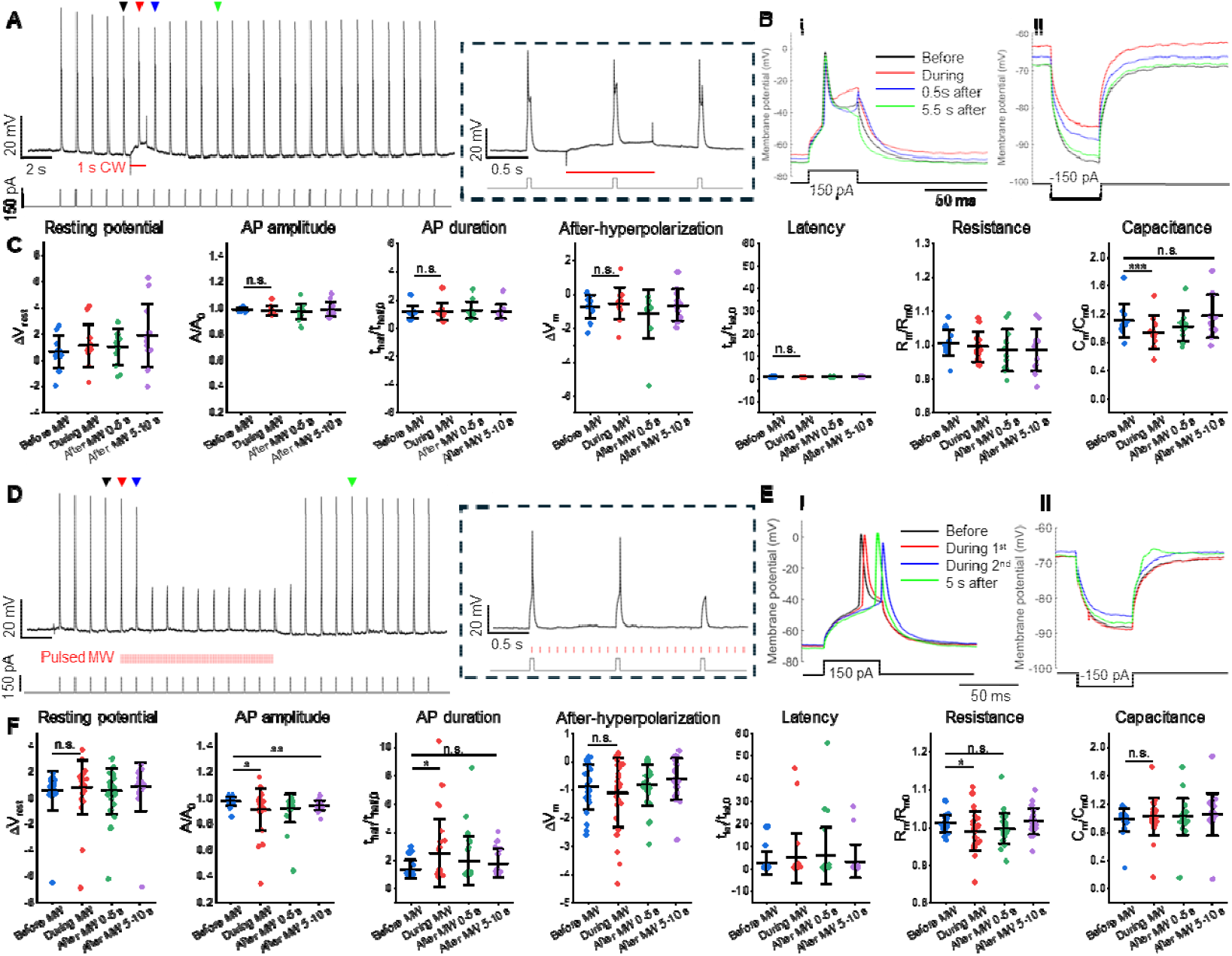
Patch clamp reveals distinct neuromodulation mechanisms. **(A)** Membrane potential of a neuron nea the MRA tip (top) in response to the current injection protocol and 1 s CW microwave (red line); inset image shows zoomed in trace during microwave exposure; **(B)** Membrane potential in response to positive (i) and negative (ii) current injections for 1 s CW microwave; **(C)** Summary of neuron membrane and action potential features for 1 s CW microwave (N = 11 neurons); **(D)** Membrane potential of a neuron near the MRA tip (top) in response to th current injection protocol and 10 s PW microwave; inset image shows zoomed in trace during microwav exposure; **(E)** Membrane potential in response to positive (i) and negative (ii) current injections for 10 s PW microwave; **(F)** Summary of neuron membrane and action potential features for 10 s PW microwave (N = 27 neurons); Significance calculated using pair-sample t test *p<0.05, **p<0.01, *p<0.001, n.s. not significant.

Next, the neurons were exposed to 0.25-0.66 W/cm^2^ 10 s PW microwave (**Figure 4D**). During the beginning of the microwave period, AP latency markedly increased until the APs were completely suppressed (**Figure 4E**). This change was not accompanied by a significant membrane depolarization. This suggests a leaky K^+^ current may be involved. Very weak correlation was found between A/A_0_ and ΔV_rest_ (**Fig. S11B**), meaning the mechanism is different than the 1 s CW case. Other notable changes include an increase in AP duration and a larger after-hyperpolarization (**Figure 4F**). When negative current was injected into the neuron, again, the membrane resistance decreased, suggesting the presence of an unspecific cationic current. These results support the hypothesis that two distinct mechanisms – thermal and nonthermal – are responsible for the differential effect of microwaves on neuronal activity.

## Discussion

In this study, we developed a MRA that can wirelessly couple and amplify microwave fields to study the bioeffects of microwaves at low operational power densities. This study is timely and relevant as the public is increasingly exposed to microwaves, particularly 0.9 – 3 GHz, via cellphones, Wi-Fi, and other devices^19^. The current standards for safe microwave exposure are primarily based on the thermal effects of microwaves which occur at higher powers^22^. While several studies have reported thermal and nonthermal neural effects of microwaves^16–19^, almost nothing is known about the acute cellular mechanisms by which they occur.

Previously, there has been some debate around whether microwaves have an excitatory or inhibitory effect on neurons, if any at all^16–19,28–30^. Our study provides new insights into the differential effects of microwave neuromodulation and their distinct cellular mechanisms. When 1 s CW microwave is applied, neurons are depolarized. When the microwave is pulsed at 10 Hz with 10% duty cycle, an equivalent dosage to the CW condition, neurons are inhibited. We observed that microwave-induced depolarization is likely caused by a rapid temperature increase (dT/dt), while microwave inhibition occurs via a nonthermal mechanism. This result is significant because inhibition occurs at lower microwave intensities than excitation, as opposed to other neuromodulation methods for which the opposite is true^6,26,31^.

We further investigated the cellular mechanisms of microwave neuromodulation by integrating the MRA into a patch clamp system. We determined that microwave-induced depolarization likely occurs via thermal activation of TRP channels, as confirmed via channel blocker experiments. Microwave-induced depolarization is therefore similar to conventional thermal stimulation techniques^26,32^. More interestingly, we discovered that nonthermal microwave inhibition is independent of membrane depolarization and occurs via a distinct mechanism. Or results suggest that leaky K+ currents are likely involved.

These results raise an important question: if the microwave inhibition is nonthermal, then how does it perturb the neural activity? We hypothesize that microwaves perturb biological systems at the molecular level. Microwave frequencies overlap with the rotational spectrum of water^33^. The orientation of water molecules at the neuronal membrane, around ions, and within ion channels has appreciable impact on neuron function^34–39^. Rotational excitation of water molecules may perturb these functions, altering membrane potential, ion solvation, and ion channel conductance. These mechanisms warrant further study to elucidate the bioeffects of microwaves and better inform exposure standards.

Our demonstration that microwaves have nonthermal, reversible effects at lower powers presents a unique method for neuromodulation therapy. Electrical stimulation, the clinical standard for neuromodulation, is highly invasive. Furthermore, electrical inhibition is achieved via high frequency stimulation which induces a depolarization block^6,40^. When used to treat conditions such as chronic pain, electrical stimulation has been reported to induce AP at the onset of the stimulus^41^, likely because depolarization block occurs with extended depolarization. Extraneous stimulation may also occur if the threshold for a depolarization block is not reached. Additionally, since electrical inhibition requires higher intensity/frequency, it has a higher potential to exceed safety limits^42,43^. Microwave neural inhibition can avoid these issues, since it does not occur via depolarization block and requires lower powers with minimal heating. The MRA occupies a volume of ~2 mm^3^ and can wirelessly couple with an external power source, making it much less invasive than electrical neuromodulation. We previously demonstrated seizure suppression in a mouse model using a microwave resonator^23^. The MRA therefore has great potential for therapeutic neuromodulation.

Overall, this study has major implications for public safety and the treatment of neurological conditions. Further study is required to elucidate the molecular mechanisms and enhance our understanding of microwave bioeffects. This knowledge will help to improve microwave exposure standards. It can also be leveraged to design highly effective and minimally invasive microwave neuromodulation devices.

## Materials and Methods

### Titanium microwave rod antenna (MRA) fabrication

The MRA was fabricated from Grade 1 pure titanium wire (Nexmetal Corporation) with diameter 0.5 mm. The wire was cut to 7.5 mm in length and the ends were polished flat using sandpaper.

### Numerical simulation

Simulations were performed in COMSOL Multiphysics 5.3a. All MRAs were modeled in bulk water medium with electrical conductivity 5.5×10-6 S/m and a relative permittivity with real part 80 and imaginary part 10. Material parameters were taken from the solid, not oxidized titanium model in the COMSOL materials library. The physical field was simulated using the Electromagnetic Waves, Frequency Domain module. The input power was 1.0 W. The MRA was oriented in the z-plane. The microwave originated from a 50 cm^2^ port with a plane wave input that has **E** polarized in the z-direction. **H** was polarized perpendicular to the MRA plane in the y-direction. Scattering conditions were used at the boundaries of the simulated area.

### Cell culture

Primary cortical neurons were harvested from Sprague-Dawley rats at embryonic day 18 (E18). Cortices were dissected from rats of either sex and digested with papain (0.5 mg/mL in Earle’s balanced salt solution) (Thermofisher Scientific). Neurons were plated onto poly-D-lysine coated glass bottom culture dishes in Dulbecco’s Modified Eagle Medium (Thermofisher Scientific) with 10% fetal bovine serum (Thermofisher Scientific). After 24 hours, medium was replaced with feeding medium consisting of Neurobasal medium supplemented with 2% B-27 (Thermofisher Scientific), 1% N2, and 1% GlutaMAXTM (Thermofisher Scientific). Fresh feeding medium was added to the culture every 3-4 days. Experiments were performed on days 10-14.

### Calcium imaging

Calcium imaging was performed on a lab-built microscope based on an Olympus IX71 microscope frame with a 10x (UPLAN FLN 10x, 0.3 NA, Olympus) or 20x air objective (UPLSAPO20X, 0.75 NA, Olympus). The sample was illuminated by a 470 nm LED (M470L2, Thorlabs), with an emission filter (FBH520-40, Thorlabs), an excitation filter (MF469-35, Thorlabs), and a dichroic mirror (DMLP505R, Thorlabs). A scientific CMOS camera (Zyla 5.5, Andor) was used to collect images at 20 frames per second. One hour before calcium imaging, cells were incubated at 37°C with 2 μM Oregon Green 488 BAPTA-1 dye (Invitrogen) for 30 minutes. The medium was then replaced with fresh medium, and cells were incubated at 37°C for another 30 minutes.

### Thermal imaging

The MRA was placed in a plastic dish and immersed in shallow DI water. The microwave waveguide was oriented with **E** parallel to the MRA plane. Microwave was delivered at the resonant frequency. Imaging was performed using a thermal camera (A325sc, FLIR). Video was captured at a frame rate of 30 Hz or 60 Hz.

### Optical temperature measurement

Precise thermal measurements were taken using an optical fiber temperature sensor (OpSens OTG-M220). The fiber tip was positioned toughing the MRA tip. Measurements were taken at a rate of 50 Hz.

### In vitro thermal controls

To fabricate the carbon coated optical fiber, the tip of a polished multimode optical fiber with a 200 μm diameter (200EMT, Thorlabs) was placed into the center of a flame for 3-5 s. In this way, a thin layer of candle soot was coated on the tip of the fiber to serve as a layer of absorber. Next, in order to stabilize the coating, a layer of PDMS was added. To prepare the PDMS, the silicone elastomer (Sylgard 184, Dow Corning Corporation, USA) was carefully dispensed into a container to minimize air entrapment. Then, the curing agent was added for a weight ratio of ten to one (silicone elastomer to curing agent). A nanoinjector deposited the PDMS onto the tip of the candle-soot coated fiber. The position of the fiber and the nanoinjector were both controlled by 3D manipulators for precise alignment, and the PDMS coating process was monitored under a lab-made microscope in real time. The coated fiber was stored overnight in a temperature-controlled environment (201C) for 12 h, to allow the PDMS to cure. A continuous 1064 nm laser was coupled into the optical fiber to heat up the carbon tip. The laser was applied for either 1 s continuously or pulsed at 10 Hz with a 10% duty cycle for 10 s. Laser pulsing was achieved using an optical chopper (Thorlabs).

### Microwave irradiation

Microwave was generated using a microwave signal generator (9 kHz to 3 GHz, SMB100A, Rohde & Schwarz) connected to a solid-state power amplifier (ZHL-100W-242+, Mini Circuits) to amplify the microwave by 51 dBm to 100 W peak power. Microwave was delivered from a 60.5 cm^2^ rectangular aluminum waveguide (WR430, Pasternack) oriented with **E** field parallel to the MRA plane. Microwave was delivered for either 1 s continuous or 10 s pulsed at 10 Hz with 10% duty cycle. Pulse modulation was achieved using a function generator (33220A, Agilent). In calcium fluorescence experiments, the MRA was placed directly on the cells. The waveguide was placed over the dish, covering it completely from above. In electrophysiology experiments, the MRA was held in the bath solution over the neurons using plastic tweezers. A QUAD 4-axis motorized micromanipulator (Sutter Instrument, Novato, CA, USA) was used to control distance to the neurons and the microwave was delivered from below the dish.

### Electrophysiology and data analysis

Both current-clamp and voltage-clamp recordings were performed using a MultiClamp 700B amplifier (Molecular Devices, San Jose, CA, USA) and digitized with an Axon Digidata 1550 (Molecular Devices, San Jose, CA, USA). Signals were lowpass filtered at 4 kHz and sampled at 20 kHz. All recordings were performed at room temperature. Recordings were conducted on cultured neurons at DIV 10–17, held at −70 mV in an external solution containing 140 mM NaCl, 3 mM KCl, 1.5 mM MgCl_2_, 2.5 mM CaCl_2_, 11 mM glucose, and 10 mM HEPES (pH 7.4). Recording electrodes were filled with a K^+^-based internal solution (135 mM K^+^-gluconate, 5 mM NaCl, 2 mM MgCl_2_, 10 mM HEPES, 0.6 mM EGTA, 4 mM Mg^2+^-GTP, and 0.4 mM Na^+^-ATP) with a resistance of 5–10 MΩ. For nucleated outside-out patch recordings, whole-cell configuration was first established. In blocker experiments, 1 nM TTX or 5 nM Ruthenium Red was applied. Data were analyzed and visualized using MATLAB (MathWorks, Natick, MA, USA). AP parameters and membrane properties were analyzed only in neurons with resting membrane potentials between −60 and −70 mV and AP amplitudes greater than 40 mV. AP amplitude was measured as the difference between the average membrane potential 1 ms before current injection and the peak of the AP. After-hyperpolarization was determined by identifying the minimum membrane potential within 100 ms following an AP peak. AP latency was calculated based on the threshold potential, defined as the membrane potential at which dV/dt first exceeded 5 V/s.

### Fluorescence data analysis

Calcium images were processed using ImageJ. The somata of all neurons with activity in a field of view near the MRA tip were selected for fluorescence measurement. Background fluorescence was removed from each frame by applying a Gaussian blur filter (sigma = 20) and subtracting it from the raw image. Traces were analyzed in Python 3.11 using a custom script. First, each trace was normalized to ΔF/F and filtered with a 40 Hz lowpass filter. The baseline of the trace was then calculated to correct for any fluctuations due to photobleaching or thermal artifacts. The baseline was calculated using an asymmetric least square smoothing algorithm with lambda = 1000 and p = 0.001. The baseline was subtracted from the filtered signal. Neural activity was quantified as the area under the curve (AUC) of the baseline-corrected trace. Plots were made using OriginPro 2021.

### Statistical analysis

Statistical analyses were performed using OriginLab 2021. Data are expressed as the mean ± SD unless otherwise stated. We used paired sample t-test for within-group comparisons and two sample t-test for between-group comparisons.

