## Supplemental figures for "Bi-modal microwave neuromodulation via thermal and nonthermal mechanisms"

**Carolyn Marar et al.**

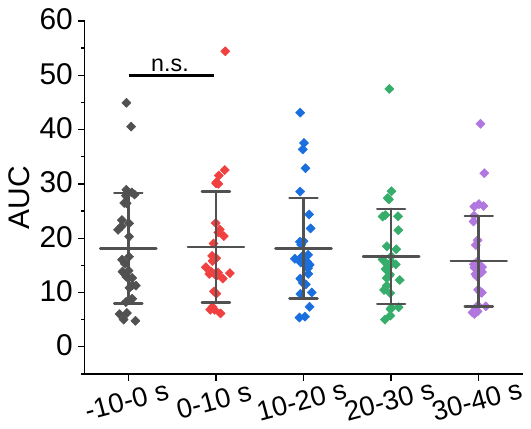

**Fig. S1:** Area under the curve of calcium fluorescence traces for cells under direct 0.66 W/cm^2^ pulsed microwave.

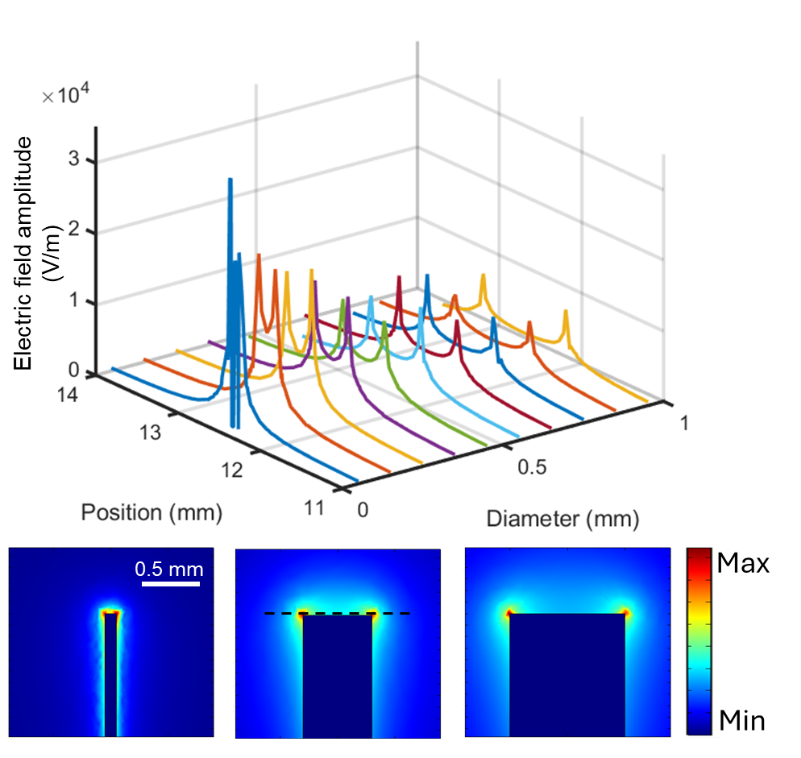

**Fig. S2:** Profile of electric field amplitude at the RMA tip (dashed line) for varying diameters

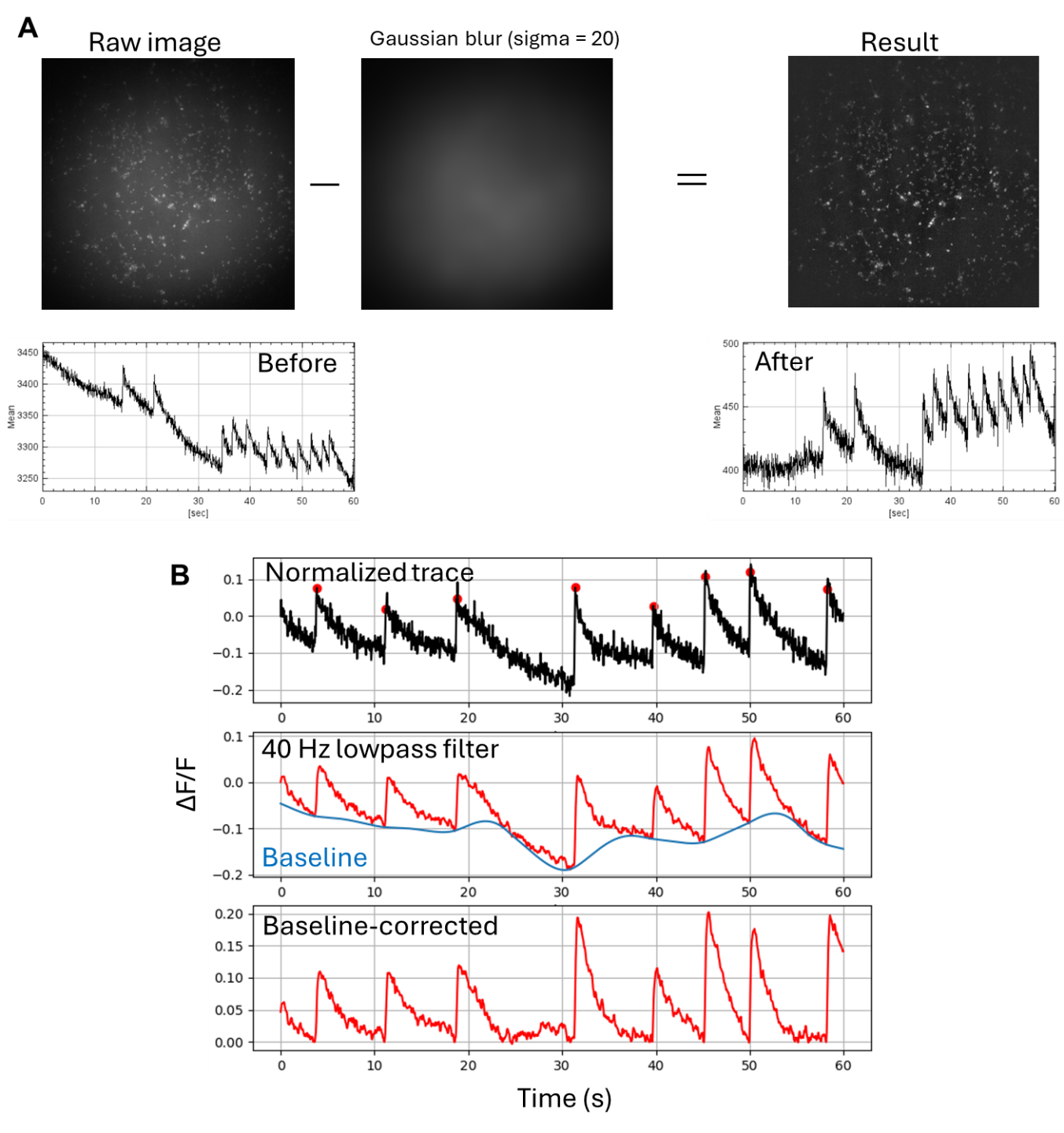

**Fig. S3: Example of data processing pipeline using ImageJ and Python**. (A) the background of the image was removed by subtracting a gaussian blur of the image; (B) Traces were normalized, lowpass filtered at 40 Hz, and baseline corrected before plotting in a heatmap.

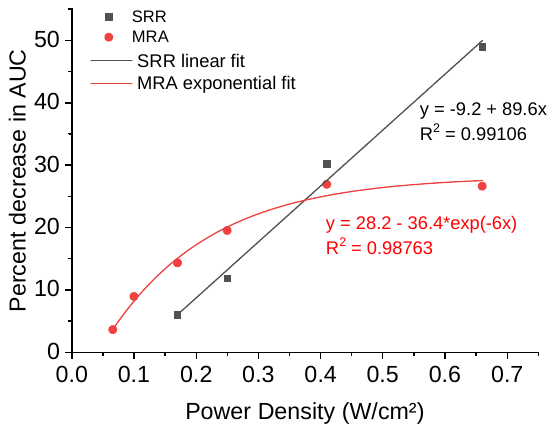

**Fig. S4:** Inhibition efficacy comparison between microwave SRR and MRA.

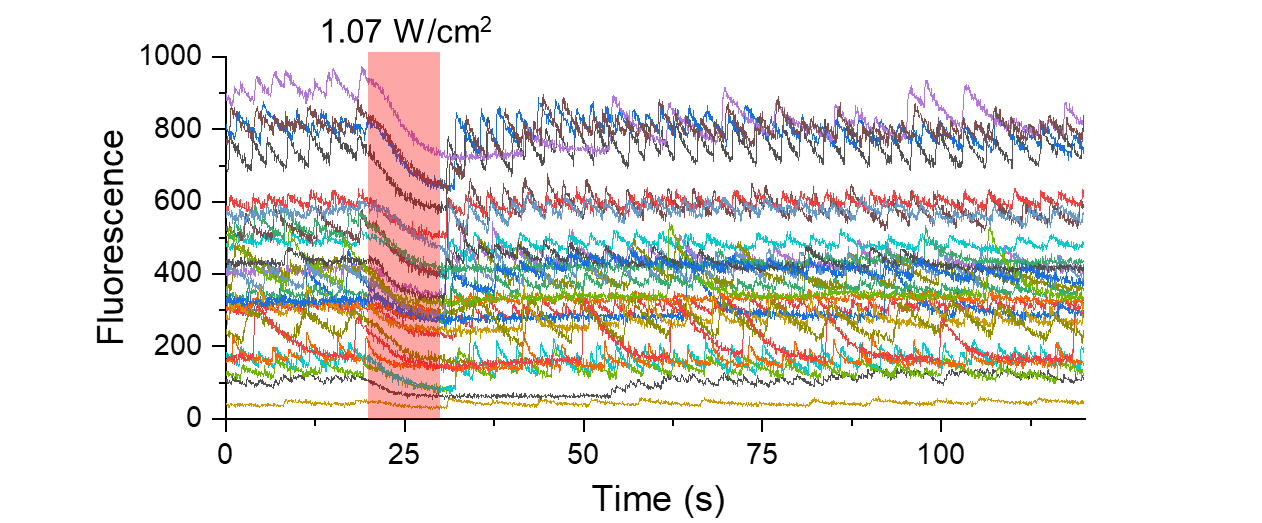

**Fig. S5:** Extended calcium fluorescence traces for neurons near the MRA exposed to 10s PW microwave.

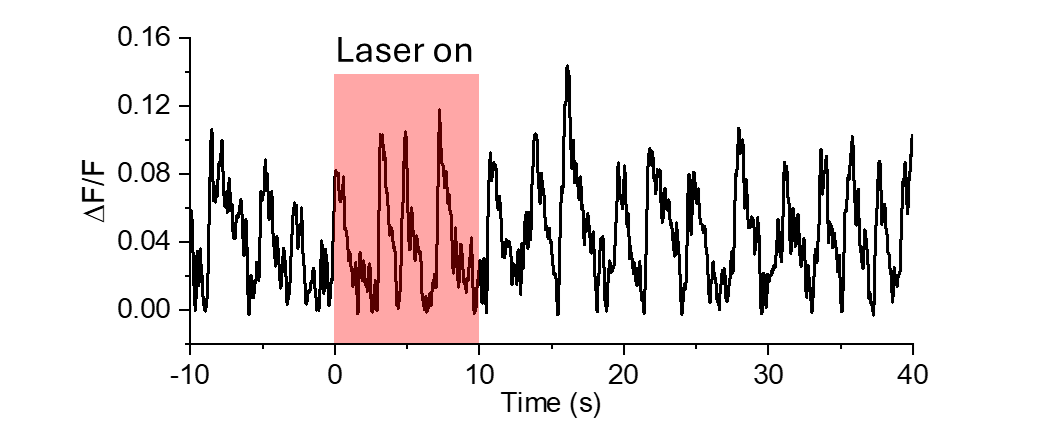

**Fig. S6:** Example thermal inhibition control trace.

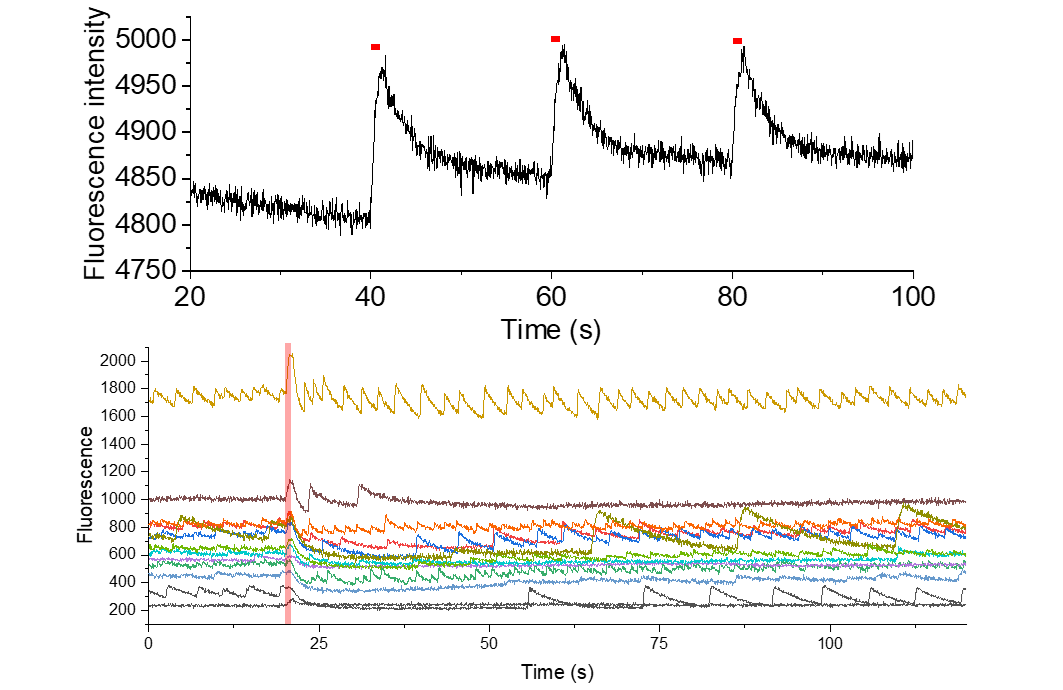

**Fig. S7:** Calcium fluorescence traces showing that 1 s CW 0.66 W/cm^2^ depolarization is repeatable and does not damage neurons.

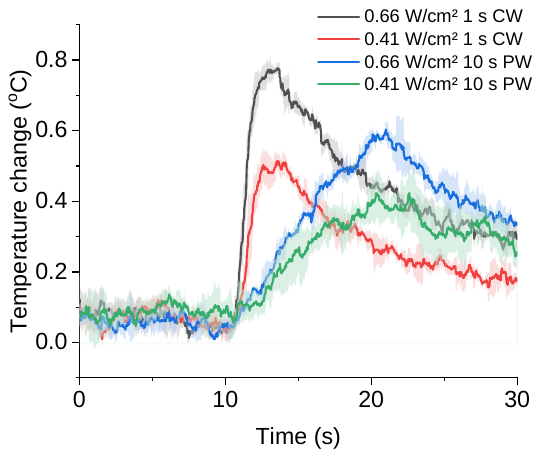

**Fig. S8:** Temperature change at MRA tip for various microwave conditions. CW: continuous wave. PW: pulsed wave.

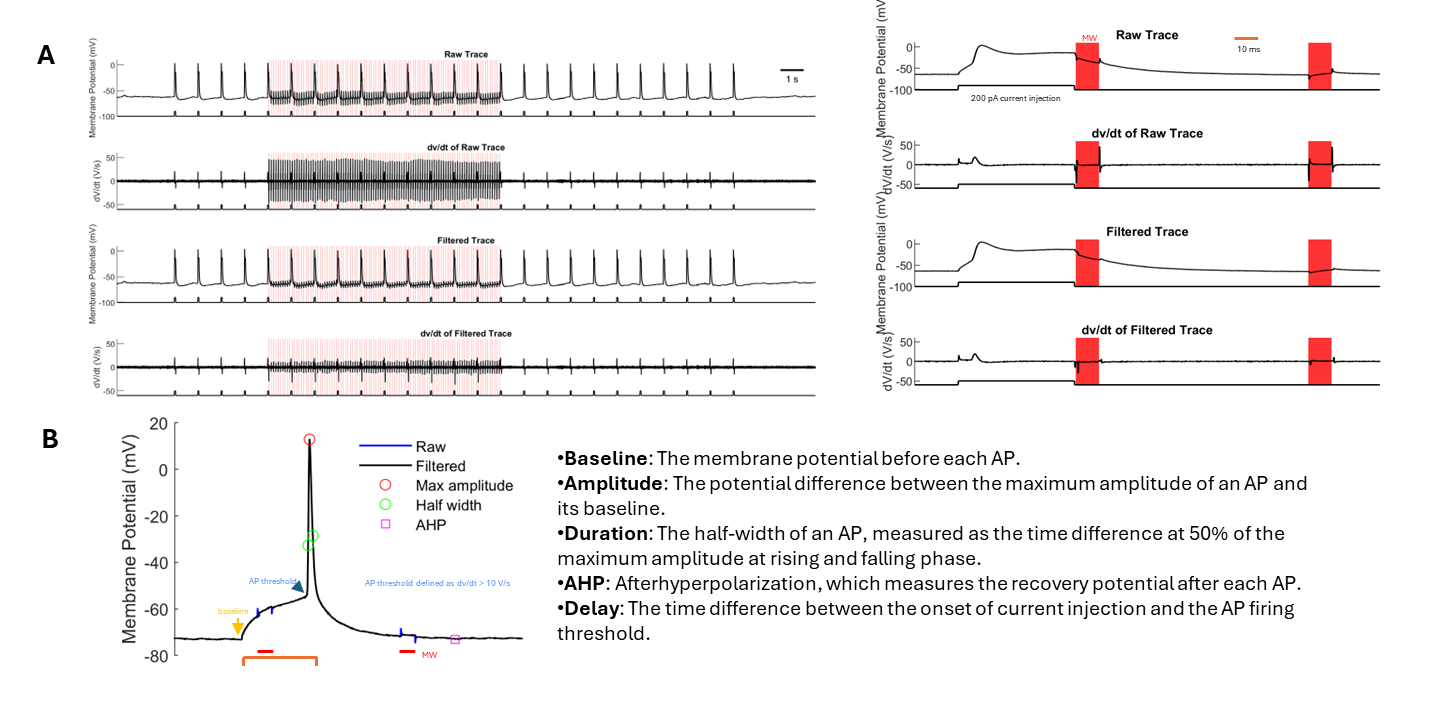

**Fig. S9: Electrophysiology processing and AP analysis. (A)** Result of bandpass filter on membrane potential traces; **(B)** Annotated AP trace showing features used for analysis.

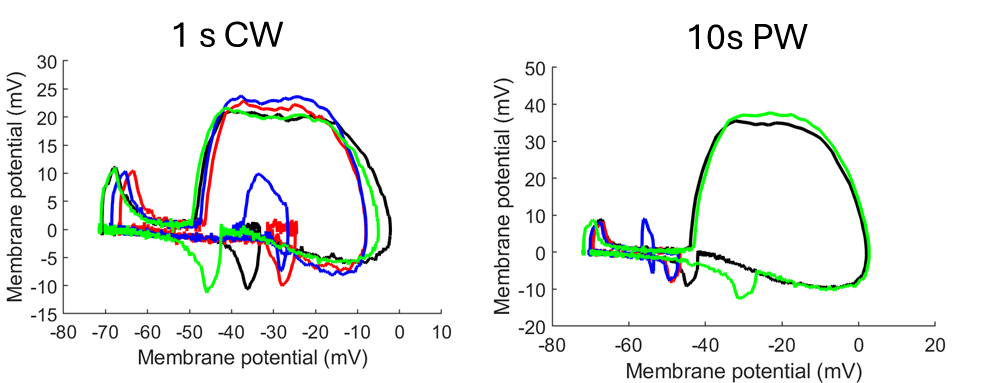

**Fig. S10.** Phase plots of the APs in Figure 4 B & E.

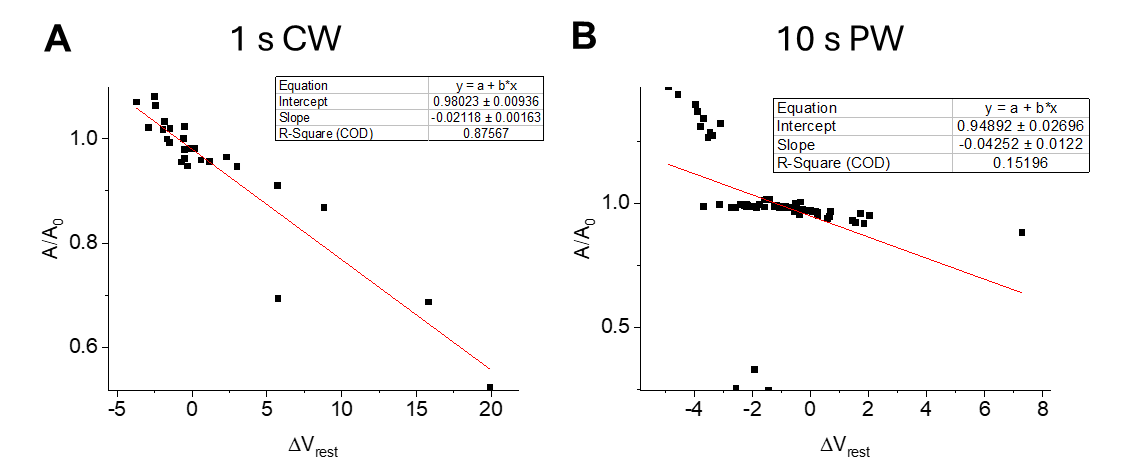

**Fig. S11:** Linear fit of AP amplitude vs change in resting membrane potentia.
